# Bacterial Ribonucleoprotein bodies maintain an acidic pH environment as a mechanism of enzyme regulation

**DOI:** 10.1101/2025.06.27.661991

**Authors:** Wade E. Schnorr, Moeka Sasazawa, Kathryn G. Dzurik, Ryan Cho, Shelby L. Millheim, Jill E. Millstone, Saumya Saurabh, W. Seth Childers

## Abstract

Phase separated biomolecular condensates create subcellular niches, yet their role in client regulation remains unclear. Here, we demonstrate that Bacterial Ribonucleoprotein bodies (BR-bodies) are acidic. Using ratiometric fluorescent probes *in vivo*, we find BR-bodies exhibit a dense-phase pH of ∼5.1, significantly lower than the near-neutral cytoplasm. Single-molecule localization microscopy and fluorescence lifetime imaging reveals that *Caulobacter crescentus* BR-bodies have spatially variable and acidic nanoscale RNase E clusters. These results question the notion of homogeneous condensates, suggesting that BR-bodies exhibit structural and biochemical diversity, which may facilitate RNA processing under stress. *In vitro*, pH gradients observed with C-SNARF-4F and RNase E CTD-pHluorin2 deteriorate with increasing buffer concentrations. Notably, the acidic microenvironment within BR-bodies enhances PNPase activity, highlighting the significance of condensate pH regulation. These findings suggest that pH modulation is intrinsic to condensates, directly influencing biochemical reactions and offering a new strategy for designing pH-sensitive drugs to target enzymes within condensates.

---

Biomolecular condensates are dynamic, membrane-less assemblies of proteins, nucleic acids, and other biomolecules that organize and regulate cellular biochemistry^1–3^. Unlike membrane-bound organelles, biomolecular condensates rely on phase separation to form distinct sub-cellular organelle-like structures that diverge in composition and density from their cytoplasmic surroundings^4–9^. Diverse biochemical pathways are regulated by biomolecular condensates in eukaryotic systems, and recent discoveries have revealed their presence and importance in bacteria, as well^1,10,11^. For example, PopZ condensates in *Caulobacter crescentus* localize at cell poles to regulate asymmetric cell division, ensuring the differential allocation of cell fate-determining proteins^12–14^. Similarly, ParB condensates are vital for chromosome segregation during cell division in several bacteria by binding and spreading along the parS sequence^15,16^. Similarly, McdB phase separation helps position carboxysomes in proteobacteria and cyanobacteria^17^. Bacteria also form ribonucleoprotein bodies (BR-bodies), analogous to eukaryotic P-bodies and stress granules, via RNase E phase separation. These condensates mediate global RNA decay by colocalizing enzymes involved in multi-step degradation, ensuring efficient turnover and preventing the accumulation of decay intermediates. Together, these condensates illustrate recent examples of how phase separation regulates essential physiology in bacteria^1,10,18^.

A major question in biology is how cells can use their 3-dimensional layout to spatially regulate the functions of enzymes and biochemical pathways. Condensates can be regulated through mass action effects, as well as unique chemical environments that influence protein folding and function. For example, in eukaryotes, membrane-bound organelles create oxidizing conditions in mitochondria and acidic conditions in lysosomes, influencing enzymatic function. Microbes generally lack membrane-bound organelles, raising the question about their capacity to generate sub-cellular niches that regulate chemical conditions like pH. The ability of condensates to generate unique chemical environments that diverge from the cytoplasm can lead to new ways to regulate enzyme functions. Here, we consider whether the recently discovered biomolecular condensates in bacteria^1,10,11^ can locally control pH to regulate function.

Weak multivalent interactions mediate phase separation, often involving intrinsically disordered regions and specific patterns of charged, polar, or hydrophobic residues^19–21^. In the “sticker and spacer” model, multivalent “stickers” promote assembly while flexible “spacers” influence solubility and dynamics^22–26^. By concentrating reactants, condensates can accelerate reactions, alter enzyme specificity, or even inhibit enzymes by sequestering substrates^27–33^.

Recent studies have identified several mechanisms by which condensates regulate pH without a membrane barrier. First, charged acidic residues (e.g. glutamate- and aspartate-rich stretches) can enrich protons within condensates, lowering local pH as seen in nucleolar condensates and RNA-binding proteins^34,35^. Second, phase separation can alter ion distribution, generating ion fluxes and interfacial electric fields as a result of variably partitioned protons and other ions, changing buffering capacity and pH^36–38^. Third, localized enzymatic activity such as ammonia production from urease-driven urea hydrolysis elevates condensate pH^39^. Fourth, condensates affect water distribution and solvation zones available for buffering^40,41^. Collectively, these mechanisms enable condensates to establish distinct pH-controlled microenvironments.

The RNase E C-terminal domain (RNase E CTD) features an intrinsically disordered region (IDR) with distinctive alternating clusters of negatively and positively charged residues (Fig. 1A), resulting in a high net negative charge and a low isoelectric point (pI) that is conserved across RNase E homologs in Alphaproteobacteria (Fig. 1C). This conservation suggests that the negative charge is crucial for BR-body function. Notably, most BR-body client proteins also exhibit a net negative charge (median = -6) (Fig. 1C). We hypothesize that ionizable side chains within BR-bodies act as intrinsic buffers, stabilizing pH to regulate enzymatic activity. Here, we provide evidence that RNase E condensates maintain an acidic environment that modulates the activity of a key client, PNPase.

**Figure 1:**
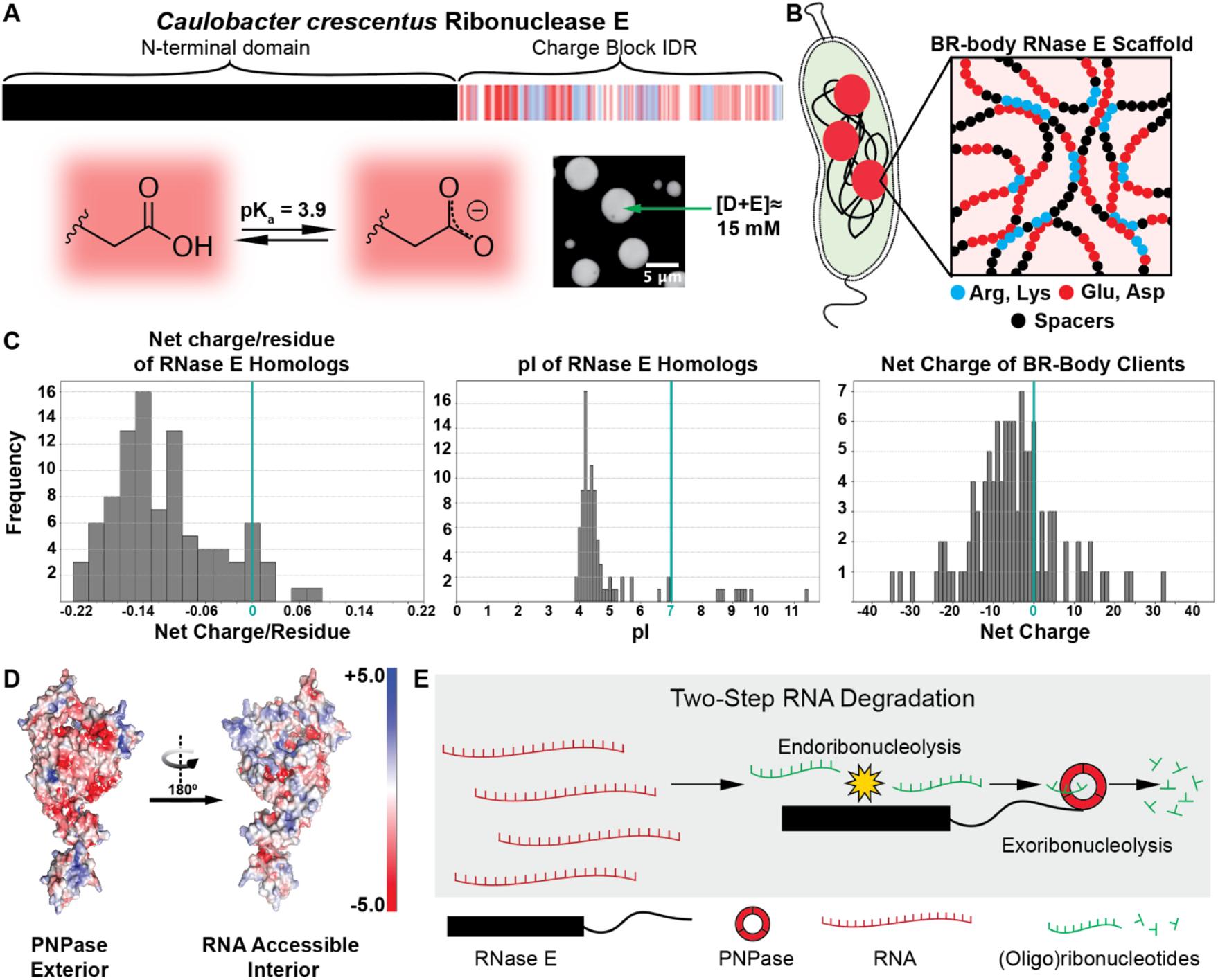
The BR-body chemical environment is defined by a high density of scaffolding and client proteins with a high density of negative charge. (A) RNase E consists of an N-terminal ribonuclease domain and a C-terminal intrinsically disordered region (IDR) with alternating clusters of negatively and positively charged amino acids. An example side chain of aspartic acid is shown in its protonated and deprotonated states to highlight the buffering ability of both aspartic and glutamic acid given the ∼15 mM concentration present in the adjacent *in vitro* RNase E-CTD condensates. (B) The cartoon representation of BR-body structure emphasizes the high net negative charge within the condensate, which may play a role in pH regulation. The zoomed-in portion is shown with other client proteins and RNA omitted for clarity. (C) Negative charge is a conserved feature among RNase E homologs in Alphaproteobacteria, which have a median net charge/residue of -0.12 and median pI of 4.4. The charge of known clients of BR-bodies also skews negative to a median net charge of -6.0. (D) Structural model of the PNPase client, showing the distribution of charges on the solvent-exposed surface compared to the RNA-binding surface. (E) Schematic showing the two-step RNA degradation process utilizing RNase E and its bound client PNPase.

## Results

### In vivo *pH measurements of* Caulobacter crescentus *expressing RNase E-pHluorin2*

We first measured BR-bodies pH *in vivo* by constructing an RNase E-pHluorin2 chimera under the native RNase E promoter in *Caulobacter crescentus*. A standard curve was generated using methylamine (pK_a_ = 10.7), benzoate (pK_a_ = 4.2), and either Tris-Cl or sodium acetate/acetic acid buffer at a desired pH. Methylamine and benzoate can diffuse across cell membranes in their neutral forms, whose relative proportions are dictated by the buffer’s pH, and will either capture protons or release protons, respectively, to match the extracellular pH. This method yielded a quantitative pH range of ∼4.8-7.5 (Fig. S1A). Ratiometric imaging of *C. crescentus* revealed a near-neutral pH of 7.0 ± 0.4 in the dilute phase and a markedly acidic pH of 5.1 ± 0.2 in the dense BR-bodies (Fig. 2A, 2B). Fluorescence intensity profiles confirmed that high RNase E density corresponds to lower pH (Fig. 2C, S1B), with acidic zones extending beyond the brightest puncta. Past single-molecule imaging experiments revealed a median of four RNase E clusters per cell(ranging from 1-10)^42^ with both large and small assemblies being acidic herein.

**Figure 2.**
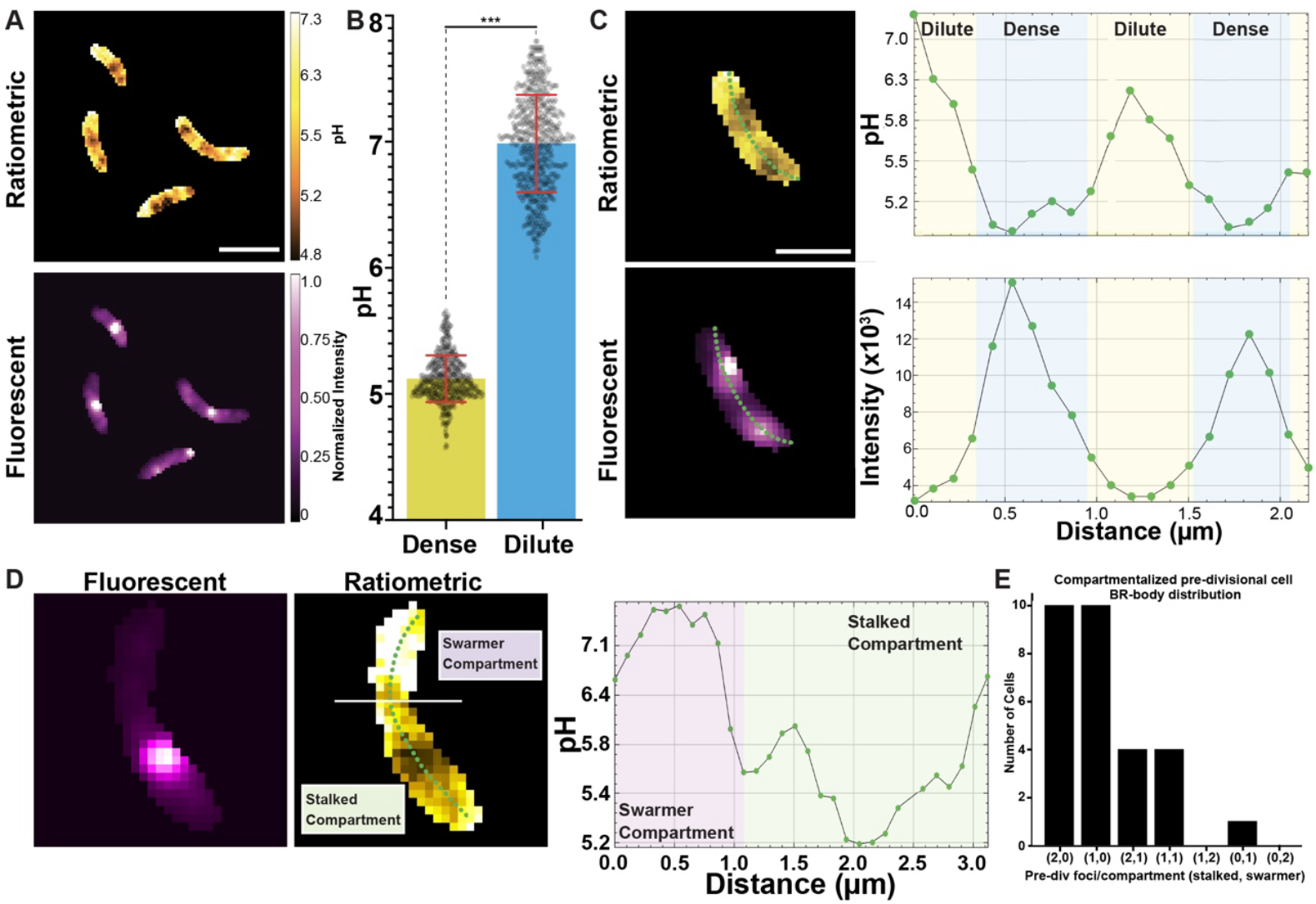
BR-bodies form distinct acidic subcellular environments in *Caulobacter crescentus* cells. (A) Ratiometric and fluorescent images of several *C. crescentus* cells endogenously expressing RNase E-pHluorin2 during mid-log phase. (B) Dense BR-bodies (n=530) and dilute phase (n=599) pH values were taken from many cells which show a dense phase pH of 5.1 ± 0.2 and a dilute phase pH of 7.0 ± 0.4 (*p<0.001*). (C) Fluorescent intensity profiles of a *C. crescentus* cell using ratiometric and fluorescent images show that high-intensity and denser BR-bodies are spatially localized to the same regions that have lower pH values. (D) Phase, fluorescent, and ratiometric images of a model pre-divisional cell and the associated ratiometric profile illustrate that swarmer compartments have fewer BR-bodies and exhibit a more neutral pH than their corresponding stalked compartments. (E) Histogram of pre-divisional cells and pairings of stalked and swarmer BR-body counts. Scale bars are 2 µm. Error bars represent standard deviations.

In *C. crescentus* pre-divisional cells, which divide asymmetrically into distinct swarmer and stalked cells, BR-bodies were more abundant in the stalked compartment, with ∼70% of cells containing one or two BR-bodies in the stalked region and none in the swarmer compartment (Fig. 2E). Consequently, the local environment around RNase E in the swarmer compartment remained more neutral, while the stalked compartment was enriched in dense-phase acidic conditions (Fig. 2D). These findings suggest that RNase E’s pH microenvironment varies upon *Caulobacter’s* asymmetric division.

### BR-bodies exhibit structural and biochemical heterogeneity

Having established that BR-bodies sustain distinct acidic microenvironments, we next explored whether these condensates possess uniform or structurally defined internal environments. Biomolecular condensates often exhibit internal heterogeneity, reflecting spatially organized functional subdomains or distinct biochemical states. We hypothesized that the acidic pH gradients identified using pHluorin2 might correspond to specialized structural subclusters within BR-bodies, potentially influencing local enzymatic activity and RNA processing. To test this idea, we utilized single-molecule localization microscopy and fluorescence lifetime imaging microscopy (FLIM) to resolve nanoscale structural organization and internal biochemical complexity in BR-bodies in live cells.

To dissect structural heterogeneities within BR-bodies, we first applied single-molecule localization microscopy^43^ using RNase E-eYFP fusion proteins (Fig. 3A). Super-resolution imaging resolved distinct sub-diffraction clusters approximately 50–100 nm in size, significantly below the diffraction limit (Fig. 3B). This fine-scale organization indicates that BR-bodies are not uniform structures but rather composed of distinct nanoscale clusters of RNase E. Such clustering could reflect localized enrichment of specific client proteins, RNA substrates, or varying stages in the RNA degradation pathway, each potentially operating within unique biochemical microenvironments.

**Figure 3.**
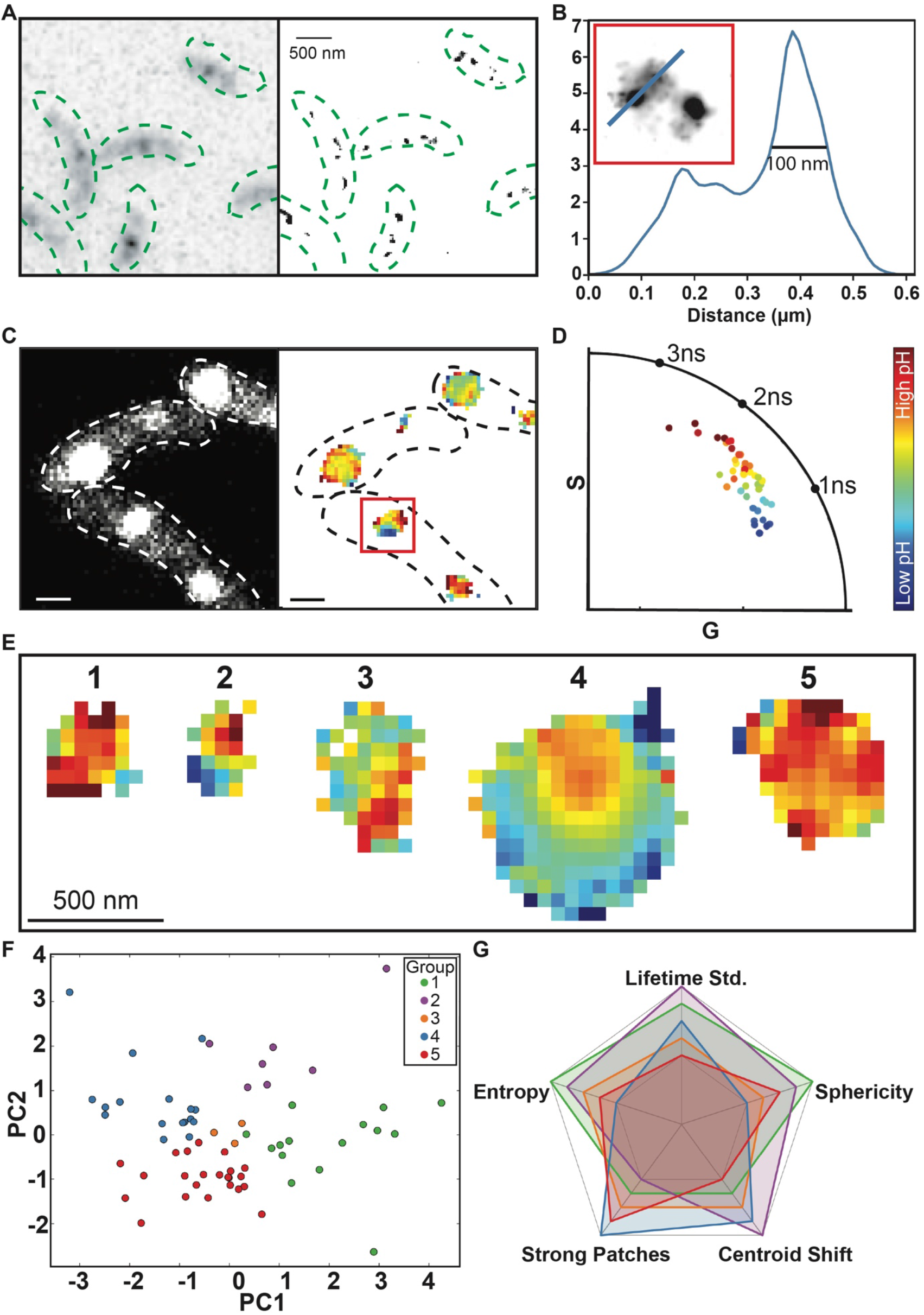
Super-resolution and fluorescence lifetime imaging (FLIM) reveal structural and biochemical heterogeneity in BR-bodies. (A) Super-resolution localization microscopy image of BR-bodies labeled with RNase E-eYFP (right) compared with the corresponding diffraction-limited image (left). (B) Resolution enhancement demonstrated by the line profile of the indicated region, showing distinct localization features separated by approximately 100 nm. (C) Intensity image (left) and corresponding fluorescence lifetime map (right) of RNase E-pHluorin2 within BR-bodies. Dashed lines outline cell boundaries. (D) Phasor plot illustrating fluorescence lifetime heterogeneity within the indicated cluster from panel (C). Colors represent lifetime values ranging from 1000 to 2300 ps. (E) Representative examples of distinct BR-body clusters categorized by their fluorescence lifetime signatures, highlighting intracluster heterogeneity. (F) Principal component analysis (left) and (G) corresponding spider plot (right) identifying critical features (Lifetime standard deviation, Sphericity, Centroid shift, Strong patches, Entropy) contributing to classification and differentiation of BR-body clusters into five distinct groups. Scale bars are 500 nm.

To explore environmental heterogeneity within these structural clusters, we employed fluorescence lifetime imaging microscopy (FLIM) with RNase E-pHluorin2^44^. FLIM revealed marked intra-cluster variability, with lifetimes ranging from ∼1 ns to ∼2.5 ns (Fig. 3C, S2A). A probability density histogram confirmed the wide distribution of lifetimes across numerous BR-bodies (Fig. S2A). Although *in vitro* calibration of purified RNase E CTD-pHluorin2 indicated fluorescence lifetimes spanning from about 1 ns at low pH (∼5) to around 3.3 ns at higher pH (∼9) (Fig. S2B), our *in vivo* measurements exhibited lifetimes predominantly below 2.5 ns, consistent with an acidic internal environment. Notably, absolute fluorescence lifetime values *in vivo* may be influenced by factors beyond pH alone, such as molecular crowding or interactions within condensates. Therefore, rather than absolute pH assignments, we emphasize the relative trends observed, where lower fluorescence lifetimes reliably indicate more acidic local environments within BR-bodies. We observed that BR-bodies within the same cell exhibited diverse fluorescence lifetimes, indicating pronounced internal biochemical complexity. Notably, the phasor plot of an individual BR-body (Fig. 3D) revealed a broad spread of lifetime components, indicating that even within a single condensate, multiple microenvironments with distinct pH states coexist. Comparison of the average fluorescence lifetimes of a cluster with the standard deviation of observed lifetimes per cluster (Fig. S2C), as well as pixel entropy (Fig. S2D) further corroborated the observations of inter- and intra-cluster heterogeneity in BR-body pH.

To systematically quantify the diversity of lifetime patterns across BR-bodies, we performed principal component analysis (PCA) and cosine similarity-based clustering on a panel of spatial features derived from each condensate. These features included lifetime variance, spatial entropy, sphericity, centroid shift, and the presence of localized high-lifetime patches. These parameters were selected to capture both the heterogeneity in the biochemical environment and the physical organization of each BR-body. This multidimensional analysis revealed five distinct groups of BR-bodies exemplified in Fig. 3E, each characterized by a unique combination of structural and lifetime features as elucidated by the PCA (Fig. 3F). To further interpret the defining characteristics of each group, we visualized the contribution of individual features using a spider plot (Fig. 3G), which revealed clear distinctions in lifetime heterogeneity and structural asymmetry across groups. Together, this classification captures a spectrum of condensate types, from compact, homogenous structures with uniform lifetimes to irregular, internally heterogeneous assemblies with sharp spatial gradients. These analyses suggest that BR-bodies exist in a range of biochemical states, potentially reflecting differences in protein and RNA client occupancy, spatial positioning within the nucleoid or near the poles, or stages of RNA degradation.

These findings collectively suggest that BR-bodies are not uniform liquid droplets but are instead heterogeneous assemblies exhibiting spatial and biochemical compartmentalization. The nanoscale clustering of RNase E, combined with internal lifetime heterogeneity and distinct condensate classifications, echoes principles observed in small-world network architectures^25^, wherein a few well-connected hubs mediate communication across a sparse but highly organized network. Just as prion-like low-complexity domains form percolated, heterogeneous networks *in vitro*, BR-bodies appear to host physically and functionally distinct subdomains that may enable parallel or staged RNA processing. This internal organization may allow BR-bodies to support diverse biochemical activities within the same condensate, possibly by tuning local connectivity, client access, or catalytic microenvironments. Such findings point to an emerging view of bacterial condensates as spatially organized, functionally modular assemblies, rather than amorphous droplets, primed for regulatory precision during stress response.

### *C-SNARF and pHluorin2 reveal an RNase E condensate pH gradient* in vitro

Past studies have utilized C-SNARF dyes and pHluorin2 to estimate the pH of polyaspartate and polyglutamate coacervates^34^, FUS^37^, BSA condensates containing urease^39^, the nucleolus^35^, and synthetic resilin- and elastin-like condensates^36^. We, therefore, compared two ratiometric pH probes to understand the pH environment within RNase E CTD condensates *in vitro*: the free small molecule C-SNARF-4F and the fluorescent protein pHluorin2. C-SNARF-4F, which exists in equilibrium between two protonation states with a pK_a_ of 6.4, has a dynamic range of ∼5-8 (Fig. S3A). An advantage of the dye is that it can be added under relatively dilute conditions to limit the quenching effects of highly crowded dye. Notably, Keating and colleagues proposed that anionic dyes entering condensates may interact with positive polyions, leading to a pK_a_ shift that affects their response to proton concentration^34^.

Compared to C-SNARF-4F, we employed an RNase E CTD-pHluorin2 chimera, which features a pH-sensitive SYG chromophore with a pK_a_ of 7.0. The β-barrel of pHluorin2 shields the chromophore from negatively charged polyions, reducing direct charge interactions with the chromophore and enhancing its sensitivity to proton concentration. Based on our *in vitro* pH titration standard curves, we found that pHluorin2 has a dynamic range of ∼6-8 (Fig. S3B).

C-SNARF-4F measurements indicated that RNase E condensates have an internal pH of 6.7 ± 0.2, compared to a dilute phase pH of 7.7 ± 0.1. Similarly, RNase E CTD-pHluorin2 condensates exhibited a dense phase pH of 6.5 ± 0.1 and a dilute phase pH of 7.6 ± 0.2. Both biosensors consistently detected a pH difference of 1.1 ± 0.2 between the dense and dilute phases (Fig. 4A, 4B). These results from two biosensors with distinct pH-sensing mechanisms collectively support that RNase E CTD condensates maintain a more acidic interior *in vitro*.

**Figure 4:**
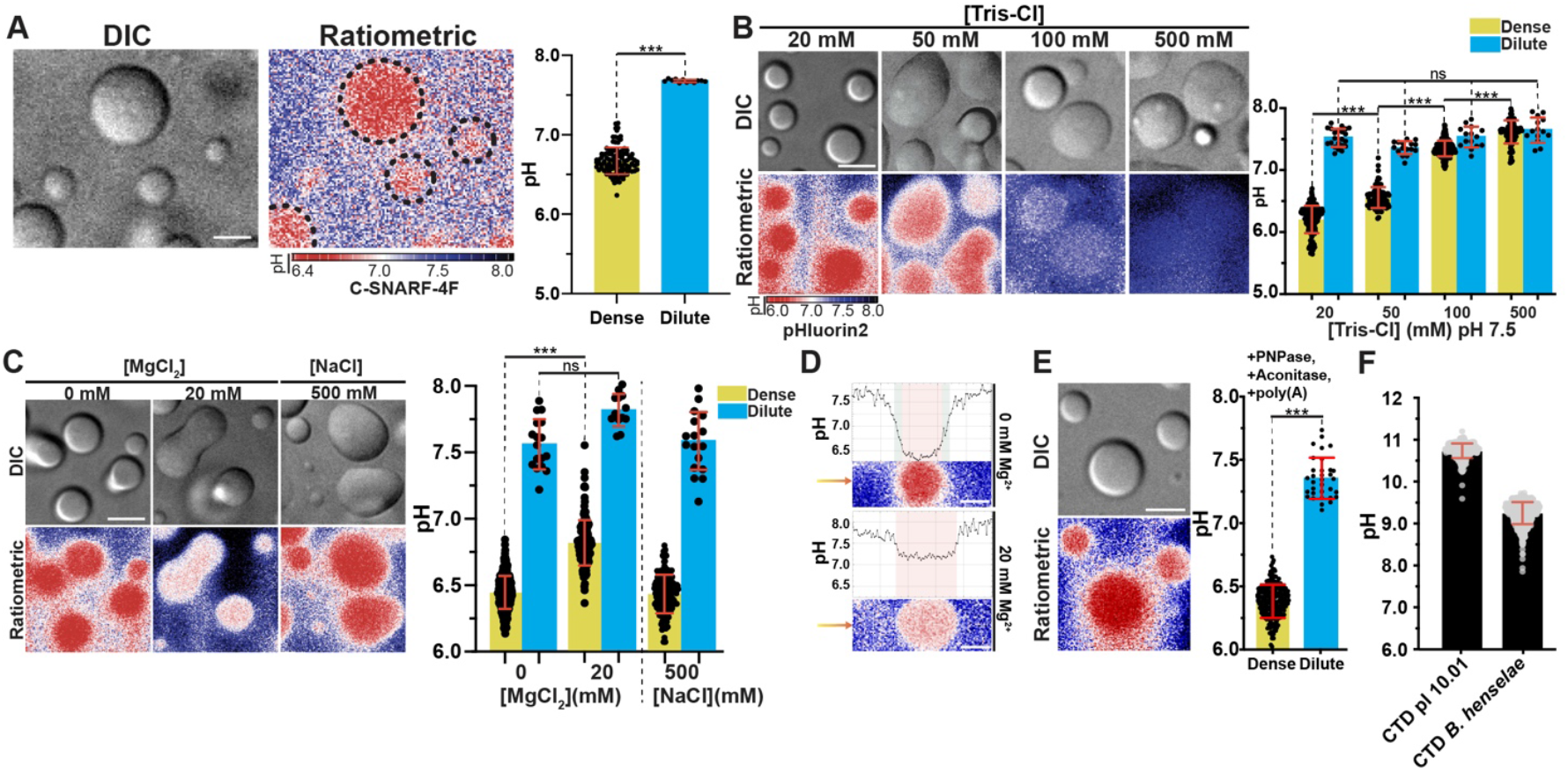
RNase E condensates act as a spatially regulated acidic pH buffer *in vitro*. (A) Representative DIC and ratiometric images of RNase E CTD condensates incubated with pH-sensitive C-SNARF-4F dye with quantitative analysis of average condensate (n=105) pH (yellow) relative to the dilute phase (cyan) (*p<0.001*). (B) Representative DIC and ratiometric images and quantitative analysis of RNase E CTD-pHlourin2 condensates (n=645) exposed to increasing concentrations of Tris-Cl buffer. Higher concentrations of the Tris-Cl buffer result in the loss of the pH gradient between the dense and dilute phases (*p<0.001*). (C) Representative images and quantitative analysis of RNase E CTD-pHluorin2 condensates (n=392) with 20 mM MgCl_2_ (n=157) and 500 mM NaCl (n=207). Adding MgCl_2_ leads to a modest increase in condensate pH from 6.5 ± 0.1 to 6.8 ± 0.2 (*p<0.001*) while adding NaCl results in a statistically insignificant change to 6.4 ± 0.2 (*p<0.001* for comparison to 100 mM NaCl). (D) Profiles of RNase E CTD-pHluorin2 with and without 20 mM MgCl_2_ show the diminished pH gradient in the presence of Mg^2+^ and a loss of a distinct zone of intermediate pH between the dense and dilute phases. (E) Representative images and quantitative analysis of RNase E CTD-pHluorin2 with PNPase, aconitase, and poly(A) (n=224). The addition of these clients does not result in a significant change in the pH of the dense and dilute phases, with values of 6.4 ± 0.1 and 7.4 ± 0.2 (*p<0.001*), respectively. (F) Quantitative analysis of the dense phase pH of mutated RNase E CTD-pHluorin2 with a pI of 10.1 and a homologous RNase E CTD-pHluorin2 from *Bartonella henselae* with a pI of 9.5 reveals dense phase pH values of 10.7 ± 0.5 (n=376) and 9.2 ± 0.5 (n=806), respectively. Scale bars are 2 µm. Error bars represent standard deviations.

Given the role of excess negative charge in other systems^34,35^, we hypothesized that the high density of aspartates and glutamates in RNase E CTD’s dense phase plays a role in buffering condensate pH. We investigated whether increasing Tris-Cl buffer concentrations, which raise the buffer-to-scaffold ratio within RNase E CTD condensates, could counteract the scaffold’s influence to neutralize the pH gradient. At Tris-Cl concentrations between 20-50 mM, the condensate maintained a lower pH of ≥ 0.80 pH units. In comparison, we observed an elimination of the condensate pH gradient at Tris-Cl buffer concentrations of 100-500 mM (Fig. 4B). This value range suggests that the RNase E CTD condensates have a buffering capacity roughly equivalent to 100 mM Tris-Cl buffer.

Next, we examined how ions regulate condensate pH because phase separation could mediate selective partitioning of ions between dense and dilute phases, leading to charge asymmetry and varied zeta potentials (Fig. S5A) as a result of induced interfacial electric potentials^38^. To test the impact of ions on the observed pH gradient, we examined the effect of adding MgCl_2_ and NaCl on RNase E CTD condensate pH. The addition of MgCl_2_ led to a statistically significant but mild increase in pH of 6.5 ± 0.1 to 6.8 ± 0.2 (Fig. 4C, S3C), indicating more specific interactions with negatively charged residues. Moreover, the larger pH gradient without MgCl_2_ displays a more gradual shift from the dilute to dense phase pH (Fig. 4D), suggesting a possible effect on the interfacial region pH as MgCl_2_ alters ionic strength and Debye length^45^. In contrast, NaCl concentrations as high as 500 mM had no statistically significant impact on the condensate pH (Fig. 4C, S3D).

To test whether ionizable side chains determine condensate pH, we engineered a synthetic *C. crescentus* RNase E CTD-pHluorin2 variant lacking many negatively charged residues but retaining known client binding sites. This construct had an estimated pI of 10.1, and we hypothesized that condensates formed in *E. coli* would exhibit a higher pH. Consistent with this, the dense phase pH was 10.7 ± 0.2. We also measured the dense phase pH of an RNase E CTD homolog from *Bartonella henselae* using a corresponding pHluorin2 chimera (estimated pI = 9.5), which yielded a pH of 9.3 ± 0.2 (Fig. 4F, S3G, S3H). These results support the idea that condensate pH is largely governed by the prevalence and identity of charged residues.

### *RNase E’s primary clients do not alter RNase E condensate pH* in vitro

Biomolecular condensates contain scaffolds that drive phase separation and client proteins that dynamically associate within the assemblies. Another key question is how clients impact RNase E condensate pH. RNase E, a scaffolding endoribonuclease essential for forming phase-separated BR-bodies, possesses more than 100 clients^46^. These clients have a range of net charges and molecular weights that could modulate condensate pH. Interestingly, over two-thirds of the clients carry a net negative charge, with a mean of -5 (Fig. 1C). Examples include three well-characterized clients that bind at near stoichiometric amounts: the exoribonuclease PNPase, the citric acid metabolic enzyme aconitase, and unstructured RNA such as poly(A). Here we examine the impact of client proteins on the buffering capacity of RNase E CTD condensates.

Without client proteins, RNase E CTD-pHluorin2 displayed an average dense and dilute phase pH of 6.5 ± 0.1 and 7.6 ± 0.2, respectively (Fig. 4B). The average condensate pH was also 6.5 ± 0.1 with 5 µM PNPase, 5 µM aconitase, or 80 ng/µL poly(A) (Fig. S3E). Titrating PNPase from 0-40 µM found statistical significance between 0-20 µM PNPase but with an effect size of 0.1-0.2 pH units, indicating large overlaps in their pH distributions (Fig S3F). With PNPase, aconitase, and poly(A) assembled simultaneously with RNase E CTD, the mean dense and dilute pH also did not change with values of 6.4 ± 0.1 and 7.4 ± 0.2, respectively (Fig. 4E). These results indicate no significant change of RNase E CTD condensate pH upon client addition *in vitro*, suggesting that the RNase E CTD condensate acidic environment is robust to alterations in its primary client composition that include changes in RNA composition during RNA degradation.

### Acidic condensate environments enhance the activity of PNPase

We considered how this robust acidic environment impacts the activity of its associated enzymatic clients. We focused on using Thioflavin T, which shifts its fluorescence (Ex=450 nm, Em=482 nm) upon binding poly(A) due to restricted rotation between its benzothiazole and benzylamine groups^47^. Its emission intensity increases with higher poly(A) concentrations and decreases as PNPase degrades poly(A), reducing binding sites (Fig. 5A); notably, when PNPase is inactive due to the absence of Mg^2+^, the emission remains unchanged (Fig. S4A).

**Figure 5:**
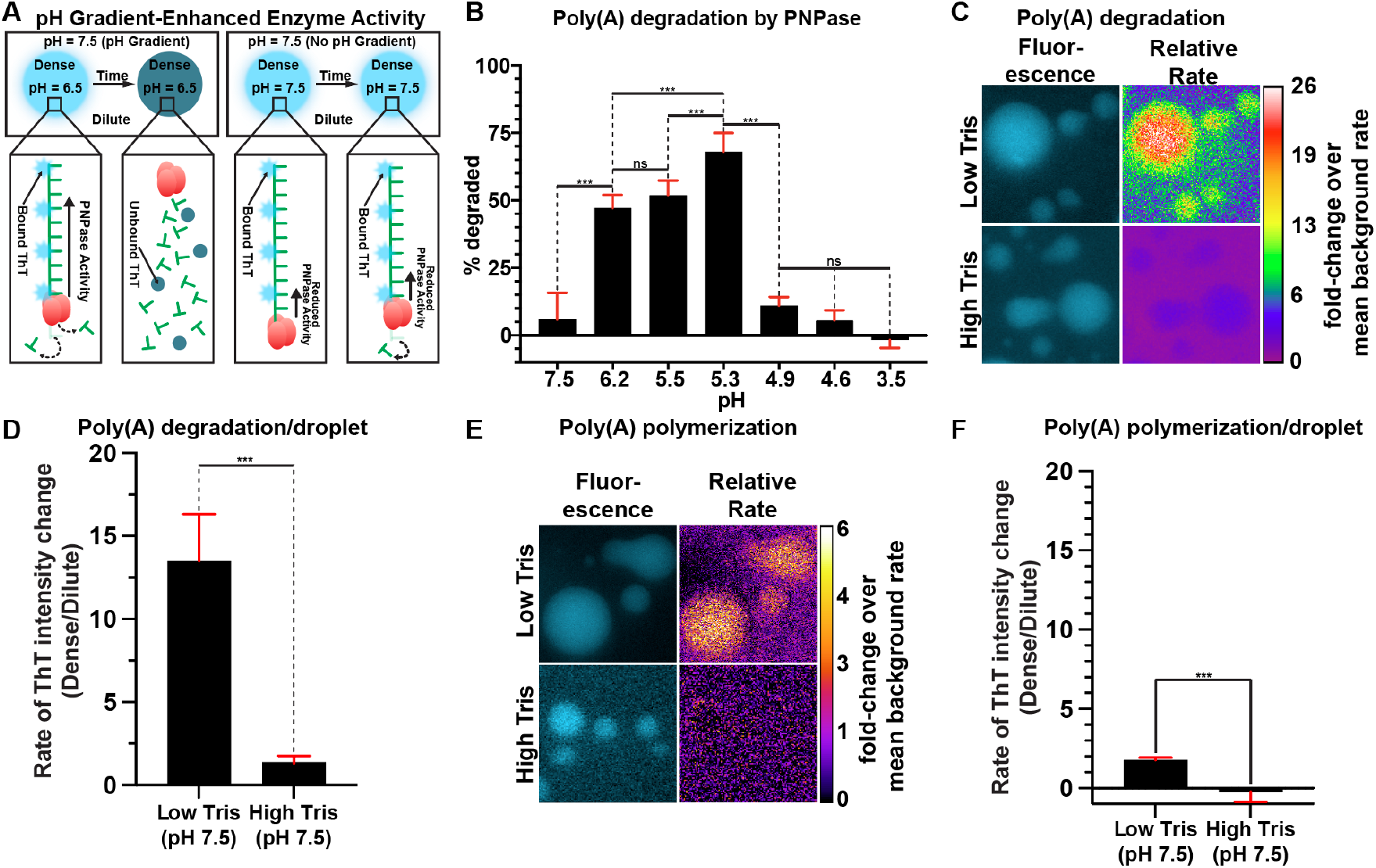
Phase separation with a pH gradient enhances poly(A) degradation and polymerization by PNPase. (A) Representation of enhanced enzyme activity due to condensate acidic microenvironment using Thioflavin T to track poly(A) degradation by PNPase. (B) The percentage of poly(A) degradation over three replicates by PNPase was measured at several pH values with a maximal activity of 68.0% ± 6.9% degraded at pH 5.3 (*p<0.001*). (C) Fluorescence imaging and relative degradation rates reveal that poly(A) degradation is strongly localized within phase-separated condensates that maintain a pH gradient (low Tris-Cl), while minimal activity is observed in the phase-separated condensates that lack a pH gradient (high Tris-Cl). (D) Analyzing the condensates (n=47) individually reveals that the relative rate of Thioflavin-T intensity change (dense/dilute ratio) shows an increased relative degradation of 13.5 ± 2.8 within phase-separated condensates that maintain a pH gradient compared to the phase-separated condensates lacking a pH gradient that show a relative degradation of 1.4 ± 0.4 (*p < 0.001*). (E) Fluorescence imaging and relative degradation rates reveal that poly(A) polymerization is strongly localized within phase-separated condensates that maintain a pH gradient (low Tris-Cl), while minimal activity is observed in the phase-separated condensates that lack a pH gradient (high Tris-Cl). (F) Analyzing the condensates (n=22) individually reveals that the relative rate of Thioflavin T intensity change (dense/dilute ratio) shows an increased relative poly(A) polymerization of 1.8 ± 0.2 within phase-separated condensates that maintain a pH gradient (low Tris-Cl) compared to the phase-separated condensates lacking a pH gradient (high Tris-Cl) that show a relative degradation of -0.2 ± 0.6 (*p < 0.001*). However, the phase-separated condensates that lack a pH gradient (high Tris-Cl) have a relative polymerization rate that is not significantly different than 0 (*p = 0.24*). Scale bars are 2 µm. Error bars represent standard deviations.

We measured fluorescence from bound Thioflavin-T:poly(A) before and after a 30-second incubation with PNPase using a bulk fluorescence plate reader assay. Applying pH-dependent standard curves (Fig. S4B), we observed PNPase activity between pH 3.5 and 7.5, with a peak at acidic levels (pH 5.3–6.2) similar to those found in BR-bodies (Fig. 2B). Specifically, poly(A) degradation was highest at pH 5.3 with 68.0% ± 6.9% degraded, but sharply decreased to 11.0% ± 3.2% at pH 4.9. At pH values above 5.3, the decrease was more gradual, with 51.8% ± 5.5% degradation at pH 5.5 and 47.1% ± 4.9% at pH 6.2 (Fig. 5B). These results suggest that the acidic environment in BR-body compartments may enhance the RNA degradation activity of PNPase.

Bulk assays reveal overall reaction dynamics but cannot distinguish activity in the alkaline dilute phase and the acidic dense phase. To address this, we used epifluorescence to monitor Thioflavin T fluorescence, which reflects poly(A) concentration changes due to PNPase activity. Under conditions in which PNPase phase separated with RNase E CTD, we calculated the average degradation rate at each pixel. By normalizing each pixel’s rate to that of the dilute background, we created a relative rate image that spatially maps PNPase activity (Fig. 5C).

Under low Tris-Cl conditions, which maintain a lower pH inside the dense condensates, the dense phase sometimes degraded poly(A) over 26-fold faster than the dilute phase, with an average 13.5 ± 2.8-fold increase. In contrast, under high Tris-Cl conditions, where the pH is equal in both phases, the increase was only 1.4 ± 0.4-fold (Fig. 5D). These results directly demonstrate that the acidic environment within RNase E condensates enhances PNPase degradation activity.

We also visualized the rate of PNPase polymerase activity, where ADP is polymerized into poly(A), using the same imaging and relative rate analysis (Fig. 5E). Starting with only ADP monomers (lacking poly(A) and inorganic phosphate), our relative rates revealed that under low Tris-Cl conditions, which maintain a lower dense phase pH, polymerization is enhanced by a factor of 1.8 ± 0.2 compared to the dilute background. Under high Tris-Cl conditions, where the pH is uniform, the rate was not significantly different from zero (Fig. 5F). These results indicate that a low pH environment, facilitated by RNase E phase separation, enhanced PNPase polymerase activity. However, RNA breakdown in the condensates remains more pronounced, supporting the role of RNase E condensates as RNA decay sites.

## Discussion

Our study demonstrates that RNase E biomolecular condensates can maintain a pH gradient *in vivo* and *in vitro*. In *Caulobacter crescentus*, BR-bodies maintain an acidic pH (∼5.1 ± 0.2) compared to the cytoplasm (∼7.0 ± 0.4) (Fig. 2A, 2B). Notably, BR-bodies are asymmetrically distributed during cell division, with more in the stalked compartment than in the swarmer (Fig. 2D, 2E). This suggests that BR-body functions may be regulated by the cell-cycle.

Super-resolution imaging and FLIM reveals that BR-bodies exhibit structural complexity, with distinct nanoscale clusters of RNase E that maintain heterogeneous acidic microenvironments rather than uniform internal conditions (Fig. 3A-3D). The distribution of fluorescence lifetimes observed within single BR-bodies demonstrates that multiple pH microenvironments are present simultaneously on a sub-condensate scale (Fig. 3E), suggesting the presence of functionally specialized subdomains. Principal component analysis further classified BR-bodies into five groups with diverse structural and lifetime features. These groups ranged from compact and homogeneous to irregularly shaped heterogeneous condensates (Fig. 3F, 3G). These variations indicate that biomolecular condensates may adopt a variety of organizational states that may be critical for their multifunctional roles in RNA metabolism.

The RNase E scaffold, rich in acidic residues (net charge of –38), appears to buffer protons much like previously observed coacervate systems, where excess charge shifts pH (Fig. 4A, 4B). Increasing Tris-Cl concentrations progressively neutralizes this gradient, indicating that aspartate and glutamate residues serve as competing buffers with Tris-Cl (Fig. 4B). In comparison, the RNase E condensates maintain a stable acidic environment in varied concentrations of salts and in the absence and presence of clients (Fig. 4C, 4E). Similarly, the vast majority of RNase E homologs have a pI between 4-6, which suggests RNase E homologs may be a common feature of BR-bodies across bacteria. Interestingly, select RNase E homologs have pIs >7 and may exhibit more alkaline or neutral interiors (Fig. 1C, 4F).

While condensates are well known to accelerate chemical reactions through mass action effects, we also found that their pH can regulate client enzyme function. *In vitro*, within RNase E CTD condensates with an acidic interior, PNPase activity increased 13.5-fold compared to the dilute phase, whereas a neutral RNase E condensate only led to a modest 1.4-fold enhancement (Fig. 5C, 5D). This pH sensitivity may arise from changes in PNPase’s conformational or oligomeric state, RNA binding, or protonation of key active site residues.

Notably, its active site contains two aspartates, a lysine, and a histidine, critical for RNA interaction, Mg^2^+coordination, and phosphate binding^48,49^. While poly(A) polymerization from ADP is also enhanced in acidic RNase E condensates, the effect is less pronounced than that on ribonuclease activity (Fig. 5E, 5F). These findings support the idea that the acidic environment of BR-bodies promotes RNA decay. Overall findings, along with evidence from studies in *Xenopus* oocytes and other *in vitro* systems^34–39^, suggest that pH gradients are a tunable feature dependent upon the net charge and pI of the condensate. Maintaining these condensate pH gradients could be crucial for regulating enzyme activity, reshaping cellular signaling, and driving metabolic processes.

## Methods

### *Imaging* Caulobacter crescentus *expressing RNase E-pHluorin2*

*Caulobacter crescentus* expressing rne::rne-pHluorin2 was grown in M2G media to OD_600_= 0.2. A 1% agarose M2G imaging pad was prepared on a glass slide (VWR), and 2 µL of the cell suspension was placed on the pad, dried for 2 min, and covered with a glass coverslip (VWR). Cells were immediately imaged using a Nikon Eclipse Ti-E inverted microscope with a Plan-Apo-λ 100x/1.45 oil objective and 518F immersion oil (Zeiss). Samples were excited at 379 nm (protonated) and 470 nm (deprotonated), with emission collected between 505-545 nm using an Andor Ixon Ultra 897 EMCCD camera. Ratiometric images were generated in ImageJ (National Institutes of Health, Bethesda, MD, USA) by dividing the 379 nm fluorescence by the 470 nm image. Phase contrast images were used to apply masks, and a yellow LUT was used for visualization. Dense BR-bodies and dilute regions were selected and measured using the Microbej^50^ plugin.

For pH calibration, cells were treated for 25 min at 30°C with 10 mM monomethylamine, 10 mM potassium benzoate, and 50 mM buffer: Tris-Cl (pH 6.2, 7.5) or sodium acetate/acetic acid (pH 4.8, 5.6). Treated cells were placed on agarose pads buffered with 50 mM of the corresponding buffer. Imaging and processing were performed as described above.

To analyze pH distribution, a 2-pixel-wide line was drawn across representative cells in ImageJ, measuring fluorescence intensity and ratio values along the line. Data were plotted using the ImageJ measurement function.

### Super-resolution microscopy and FLIM of RNaseE-pHlourin2

*Caulobacter crescentus* cells expressing rne::rne-eYFP or rne::rne-pHluorin2 were grown overnight in M2G medium, diluted to OD600= 0.05, and incubated at 30 °C with shaking at 200 rpm. Once cultures reached OD600= 0.2, cells were imaged on a 1% agarose M2G pad.

Single molecule localization microscopy was performed on an inverted microscope (Nikon Ti2) equipped with DIC optics, an oil immersion objective (Nikon PlanApo, 100x, 1.45 NA), and a sCMOS camera (Photometrics Prime 95B) with a system magnification of 0.11 µm/ pixel. A 514 nm continuous wave laser was used for exciting RNase E-eYFP in live *Caulobacter* cells. The sample was pre-bleached at high laser power (800 W/cm^2^) to get to single molecule density followed by imaging 1000 frames at 20 ms exposure time. Data was processed using Thunderstorm plugin in Fiji.

Fluorescence lifetime imaging was performed on a Nikon Eclipse Ti2 microscope equipped with a confocal scanning system (Abberior). A 100x, NA 1.45 oil-immersion objective and a 1 Airy-unit pinhole was used. The confocal detectors were connected to a Time-Correlated Single-Photon Counting (TCSPC) module (Becker & Hickl). For the measurement of pHlourin2 lifetime, 488nm excitation pulsed diode laser (40 MHz) at 5.15% power; emission at 505–550 nm was employed. All FLIM data were acquired in line-scanning mode (pixel dwell time 10 µs) using SPCM Data Acquisition software (v9.89; Becker & Hickl), with 90 s acquisition per frame. Images were segmented by pixel intensity and analyzed for decay curves and phasor plots in SPCImage NG Data Analysis software (v8.87.0; Becker & Hickl).

### Cluster analysis

Image data was first processed through a binary mask, which utilized lifetime statistics to identify spatial heterogeneity and shape. Quantitative feature extraction was applied utilizing ‘RegionProps’ module from ‘skimage.measure’ in Python. Over 15 features were extracted, but they were selectively eliminated after data visualization via pair plots and a heatmap revealed collinearity. The goal of this pre-processing was to turn image data into a tabular CSV file with unique and impactful features for further statistical analysis. The final five features to determine heterogeneity were lifetime standard deviation, sphericity, centroid shift, number of strong patches, and entropy per pixel.

After pre-processing, the CSV data was then processed through a cosine similarity program. The cosine similarity results were then utilized as the basis of a hierarchical clustering model. This model grouped all data points on feature similarity. Principle Component Analysis (PCA) was then used to reduce the dimensionality of our data to 2D for cleaner visualization. Hierarchical clustering data was then overlaid on the PCA graph utilizing a hue. Once groups were established with hierarchical clustering and PCA, F values and P values were extracted via Python and used for further analysis regarding which traits contributed most to heterogeneity and cluster groupings.

### RNase E CTD, RNase E CTD-pHluorin2 Expression and Purification

RNase E(451-898) (RNase E CTD) and RNase E CTD-pHluorin2, both with N-terminal His_6_-MBP tags, were expressed in BL21(DE3) cells transformed with pTEV5-RNase E CTD or pTEV6-RNase E CTD-pHluorin2. Cultures were grown to OD600= ∼0.5, induced with 1 mM isopropyl-β-D-thiogalactopyranoside (IPTG), and incubated for 3 h at 37°C. Cells were harvested (4°C, 4000 × g, 10 min), resuspended in HEPES pH 8.0, 500 mM NaCl, pelleted again, and stored at -80°C.

Thawed cells were resuspended in 100 mL lysis buffer (500 mM NaCl, 20 mM Tris-Cl pH 7.4, 20 mM imidazole pH 7.0, 1 mM DTT, 0.1% Triton X-100) supplemented with 20 U Pierce™ Universal Nuclease (Thermofisher™), SIGMAFAST protease inhibitor (MilliporeSigma), and 0.1 mg/mL lysozyme. Lysis was performed via an Avestin Emulsiflex-C3 at 15,000 psi for 30 min. Debris was removed by centrifugation (4°C, 20,000 × g, 45 min).

The supernatant was purified using a 5 mL HisTrap™ FF column (Cytiva) on a Cytiva Äkta™ pure 25. After equilibration and 10 column volume washes, proteins were eluted with 20 mM Tris-Cl pH 7.4, 200 mM NaCl, 200 mM imidazole, 1 mM DTT. The eluted protein was concentrated to ∼20 mg/mL using a 50 kDa MWCO Sartorius Vivaspin® concentrator, then incubated with TEV protease (1 mg per 20 mg protein, 2 h at RT). Cleaved protein was separated via a HiPrep Sephacryl S-200 HR size exclusion column (Cytiva) equilibrated in 20 mM Tris-Cl pH 7.5, 200 mM NaCl, 1 mM DTT. Final protein fractions (∼8 mg/mL) were concentrated (30 kDa MWCO Sartorius Vivaspin®), aliquoted, and stored at -80°C.

### Imaging RNase E with C-SNARF-4F

RNase E CTD was diluted to 20 µM in a buffer containing 20 mM Tris-Cl (pH 7.5), 100 mM NaCl, 10% PEG-8000, and 10 µM C-SNARF-4F (ThermoFisher) at room temperature. Components were added sequentially: water, Tris-Cl, NaCl, RNase E, C-SNARF-4F, and PEG-8000. The solution was pipetted into a 1 mm adhesive spacer well (Electron Microscopy Sciences) on a microscope slide (VWR) and sealed with a glass coverslip (VWR). Samples were inverted and incubated for 20 minutes at room temperature, then imaged using the same microscope setup as live-cell fluorescence measurements. Excitation was at 470 nm, with emission collected at 579-631 nm (protonated state) and 669-741 nm (deprotonated state) to generate ratiometric images. Image analysis in ImageJ involved background subtraction (rolling ball radius = 50) followed by image division.

For standard curve generation, PEG-8000 was omitted, and buffers at defined pH values were used: Tris-Cl (pH 6.6, 7.1, 7.7, 8.1) or sodium acetate/acetic acid (pH 3.5, 4.0, 5.0, 5.5, 5.9). Images were acquired under identical conditions and fit using nonlinear regression (Fig. S2A) of the form:

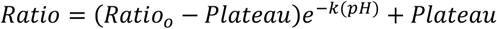

### Imaging RNase E-pHluorin2 with Tris-Cl, MgCl_2_, and NaCl titrations

For the Tris-Cl titration, RNase E CTD-pHluorin2 (20 µM) was diluted into buffers containing 20, 50, 100, or 500 mM Tris-Cl (pH 7.5), 100 mM NaCl, and 10% PEG-8000. For the MgCl_2_ titration, RNase E CTD-pHluorin2 (20 µM) was diluted into a buffer containing 20 mM Tris-Cl (pH 7.5), 100 mM NaCl, and 0, 20, 50, 75, 100, 200, or 500 mM MgCl_2_, with 10% PEG-8000. For the NaCl titration, RNase E-pHluorin2 (20 µM) was diluted into a buffer containing 20 mM Tris-Cl (pH 7.5), 100, 250, or 500 mM NaCl, and 10% PEG-8000.

All samples were prepared and imaged using the same slides, coverslips, spacers, incubation times, and imaging system as with C-SNARF-4F. Excitation was performed at 379 nm and 470 nm, with emission collected between 505-545 nm for ratiometric image generation.

### Imaging RNase E-pHluorin2 with clients

RNase E CTD-pHluorin2 (20 µM) was diluted in a buffer containing 20 mM Tris-Cl (pH 7.5), 100 mM NaCl, either 5 µM PNPase, 5 µM aconitase, or 80 ng/µL poly(A) (or combined together), and 10% PEG-8000. Components were added in the following order: water, Tris-Cl, NaCl, RNase E-pHluorin2, client(s), and PEG-8000. All solutions were imaged using the same slides, coverslips, spacers, incubation times, and imaging system as with C-SNARF-4F and the RNase E CTD-pHluorin2 titration experiments.

Samples were excited at 379 nm, corresponding to the protonated state, and 470 nm, corresponding to the deprotonated state. Emission was collected between 505-545 nm for ratiometric image generation. The final ratiometric images were created by dividing the fluorescence image captured at 379 nm by the image obtained at 470 nm (after background subtraction with a rolling ball radius of 50) using ImageJ software. A custom red-to-blue LUT was applied in ImageJ to visualize the resulting ratios.

### PNPase activity assay

PNPase activity measurements were conducted using a plate reader (Tecan Infinite M1000 Pro). Samples were prepared by diluting PNPase (2.5 µM) in a buffer containing 20 mM Tris-Cl (for pH 6.2 and 7.5) or 20 mM sodium acetate/acetic acid (for pH 3.5, 4.6, 4.9, 5.3, and 5.5), 100 mM NaCl, 20 ng/µL poly(A), 10 µM Thioflavin T, 500 µM MgCl_2_, and 4 mM phosphate. Components were added in the following order: water, buffer, NaCl, PNPase, poly(A), MgCl_2_, Thioflavin T, and phosphate.

For standard curves, each pH condition included poly(A) concentrations of 20, 15, 10, 5, and 0 ng/µL, with all other components unchanged except for the exclusion of MgCl_2_ (Fig. S4B).

Samples (70 µL) were loaded into a 384-well plate, and fluorescence intensity was measured with an excitation wavelength of 450 nm (5 nm bandwidth) and emission at 485 nm (5 nm bandwidth). Each sample received eight readings per timepoint, measured sequentially. Standard curve fits were either exponential or linear depending upon comparisons between the models.

The plate reader temperature was maintained at 26–27.5°C. For activity measurements including MgCl_2_, all components except phosphate were added to the wells, and phosphate was mixed for ∼8 seconds before initiating sequential measurements.

### *Calibration Curve for* in Vivo Caulobacter crescentus *pH Measurements*

The *C. crescentus* strain containing the rne::rne-pHluorin2 construct was grown in M2G media to OD600= 0.3. Cells were pelleted by centrifugation at 5,000 × g for 10 minutes at room temperature, and the clarified supernatant was transferred to a fresh tube. This supernatant was used to prepare calibration media by supplementing it with 10 mM monomethylamine and 10 mM potassium benzoate. To establish specific pH conditions, 50 mM Tris-Cl was added for pH 6.2 and 7.5, while 50 mM sodium acetate/acetic acid was used for pH 4.8 and 5.6.

Cells were resuspended in an equal volume of calibration media to maintain the original OD600and incubated at 30°C for 15 min. in the corresponding pH buffer. Each condition was prepared separately to ensure consistent timing.

Simultaneously, 1% agarose imaging pads were prepared, supplemented with the same 50 mM buffer used for each pH condition. Following incubation, 2 µL of the cell suspension was applied to the imaging pad, dried, and imaged.

Samples were excited at 379 nm and 470 nm, with emission collected between 505-545 nm to generate ratiometric images. The fluorescence image obtained at 379 nm was divided by that at 470 nm. Regions of cells without apparent BR-bodies were analyzed for fluorescence ratios, and these data were used to construct the calibration curve (Fig. S1A), which was fitted to the following equation:

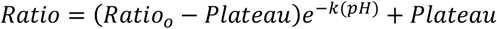

### Electrostatic Visualization of PNPase Structure

The three-dimensional structure of PNPase was obtained from the Protein Data Bank (PDB). Solvent molecules, including crystallographic water, were removed using PyMOL (Schrödinger, LLC). To determine the protonation states of titratable residues (histidine, aspartic acid, glutamic acid, lysine, and arginine) at the experimental pH, the protein structure was processed using pdb2pqr^51^, which assigned appropriate protonation states based on the PROPKA algorithm. The processed structure was saved in PQR format for further electrostatic calculations.

The electrostatic potential of the protein surface was computed using the Adaptive Poisson-Boltzmann Solver (APBS)^52^. The input PQR file was used to generate the electrostatic potential map, with the following parameters: a dielectric constant of 2 for the protein interior and 78.5 for the solvent, a solvent radius of 1.4 Å, and a grid spacing optimized to balance resolution and computational efficiency. The Poisson-Boltzmann equation was solved under physiological ionic conditions, and the resulting electrostatic potential was saved for visualization.

PyMOL was used to visualize the electrostatic potential on the protein surface. The PDB structure was loaded, followed by the DX file generated by APBS. The “APBS Electrostatics” plugin within PyMOL was utilized to map the electrostatic potential onto the molecular surface, applying a color gradient from red (negative charge) to blue (positive charge). Visualization settings, including surface transparency, color ramp adjustments, and contouring, were optimized to enhance the clarity of electrostatic distributions.

### RNase E Homolog and Client Charge

Homologs were identified by BLAST-searching the *C. crescentus* RNase E (RNE) protein sequence against the National Center for Biotechnology Information (NCBI) database^53,54^. Intrinsically disordered regions within the protein sequences were predicted using Metapredict, while amino acid composition was analyzed using ProtParam^55,56^. BR-body client proteins were identified by Nandana *et al*.^46^ Client protein sequences and amino acid composition were processed identically to the homologs.

### Designing and Imaging Synthetic RNase E CTD

The synthetic RNase E CTD (estimated pI=10.1) construct was generated by modifying the *Caulobacter crescentus* RNase E C-terminal domain (CTD). Aspartic acid (D) and glutamic acid (E) residues within three key negatively charged regions were substituted with neutral residues: E559S, E560T, D561S, D562T, D563S, D564T, D570S, D571T, E572S, E573T, E574S, E575T, D576S, D577T, D581S, D582T, E583S, D584T, D585S, E589T, D590S, D592S, D593T, D594S, D595T, E632S, E634T, E636T, E638S, D646T, D648S, D654S, D655T, E826T, D829S, E833S, E837T, E842S, E845S, E847T, E850S, E859S, E863T, E869S, E872T, E873S, E878S, D880T, E883S. These substitutions were designed to reduce the net negative charge while preserving known client-binding motifs.

### Microscopy Analysis of PNPase Activity

For poly(A) degradation, RNase E CTD (20 µM) was diluted in a buffer containing 20 mM or 100 mM Tris-Cl (pH 7.5), 100 mM NaCl, 500 µM MgCl_2_, and 10% PEG-8000. Additional components included PNPase (2.5 µM), poly(A) (20 ng/µL), Thioflavin T (10 µM), and phosphate (4 mM). Components were added sequentially in the following order: water, Tris-Cl, NaCl, RNase E, PNPase, poly(A), MgCl_2_, Thioflavin T, PEG-8000, and phosphate.

Since PNPase requires phosphate for activity, phosphate was omitted until just before imaging. Samples without phosphate were deposited on a 22 × 60 mm glass coverslip (VWR), incubated for 10 minutes in a humidity chamber containing a moist paper towel to minimize evaporation, and then placed on the microscope stage. Phosphate was added *in situ*, and images were acquired every 10 seconds with 440 nm excitation, 480 nm emission, and a 50 ms exposure time.

For poly(A) polymerization, RNase E CTD (20 µM) was diluted in a buffer containing 20 mM or 100 mM Tris-Cl (pH 7.5), 100 mM NaCl, 500 µM MgCl_2_, and 10% PEG-8000. Additional components included PNPase (2.5 µM), ADP (50 µM), and Thioflavin T (10 µM). Components were added sequentially in the following order: water, Tris-Cl, NaCl, RNase E, PNPase, ADP, Thioflavin T, PEG-8000, and MgCl_2_.

Since PNPase also requires MgCl_2_ for activity, MgCl_2_ was omitted until just before imaging. Samples without MgCl_2_ were deposited on a 22 × 60 mm glass coverslip (VWR), incubated for 10 minutes in a humidity chamber containing a moist paper towel to minimize evaporation, and then placed on the microscope stage. MgCl_2_ was added *in situ*, and images were acquired every 10 seconds with 440 nm excitation, 480 nm emission, and a 50 ms exposure time.

Relative rate images were generated by selecting frames 30 seconds apart for degradation and 10 seconds apart for polymerization. Pixel intensity differences between frames were divided by the respective time intervals to compute pixel-specific mean intensity rates of change. These rates were then normalized by dividing the mean background intensity rate of change for each pixel. Finally, the relative rates of multiple condensates were averaged to enable quantitative comparisons between low and high Tris-Cl conditions.

