## Supplemental Information for "Bacterial Ribonucleoprotein bodies maintain an acidic pH environment as a mechanism of enzyme regulation"

### Supplementary Text

#### *Ratiometric and fluorescence intensity colocalization analysis of BR-bodies*

To assess the spatial correspondence between low-pH regions in ratiometric images and high-fluorescence regions in single-channel images, we performed three colocalization analyses with progressively stricter penalties for non-overlapping pixels. In all cases, we compared the pixel locations residing within the first quartile of intensity (lowest pH) of the ratiometric image (subset *A*) to those within the fourth quartile of intensity of the single-channel fluorescence image (subset *B*), evaluating how well the lowest pH pixels corresponded to the brightest BR-bodies. The first of such analyses was the overlap coefficient—which compares the number of colocalized pixels to the smaller subset size $\left(O = \frac{|A \cap B|}{\min(|A|,|B|)}\right)$—yielded a value of 0.72 ± 0.02. Next, the Dice-Sørensen coefficient, which calculates the fraction of colocalized pixels relative to the average subset size $\left(D = \frac{2|A \cap B|}{|A|+|B|}\right)$, also produced a value of 0.72 ± 0.02, consistent with the overlap coefficient due to the similarity in sizes of subsets *A* and *B*. Finally, the Jaccard index, the strictest measure that compares the overlap to the union of both subsets $\left(J = \frac{|A \cap B|}{|A \cup B|}\right)$, resulted in a value of 0.56 ± 0.02. These values fall within the range of significant overlap in all methods that were used (Fig. S1B).

#### *Estimation of Debye length and zeta potential as functions of ionic strength*

A pH gradient establishes an interfacial electric potential, influenced by the local arrangement of proteins at the condensate interface and differential ion partitioning between phases. This potential decays exponentially over a characteristic distance known as the Debye length, which depends on the ionic strength of the solution. Thus, altering ion concentrations can modify the Debye length (Fig. S5B). The average electric field in the interfacial region can be approximated by dividing the potential drop across the interface by the thickness of the electric double layer, typically on the order of the Debye length.

The Boltzmann distribution describes how ion concentration—and thus pH—decays exponentially over a distance defined by the Debye length. Additionally, selective partitioning of specific ions into the condensate can shift the local pH by altering both buffering capacity and ionic strength, thereby modifying the equilibrium conditions predicted by the Boltzmann distribution and changing the spatial extent of pH variation across the interface.

The Debye length is given by a linearized Poisson-Boltzmann model:

$$\lambda_D = \sqrt{\frac{\varepsilon_0 \varepsilon_r k_B T}{N_A e^2 \sum_i c_i z_i^2}}$$

Because the Debye length depends on the presence of any charge species ($z_i$), proteins with titratable residues can significantly influence its value through changes in their protonation states. In some cases, both protonation state changes and selective ion partitioning contribute to Debye length variations. Consequently, zeta potential measurements, which depend in part on the Debye length, follow similar trends. Notably, even in cases where the pH gradient is eliminated, local charge distributions can still generate a nonzero zeta potential (Fig. S5A).

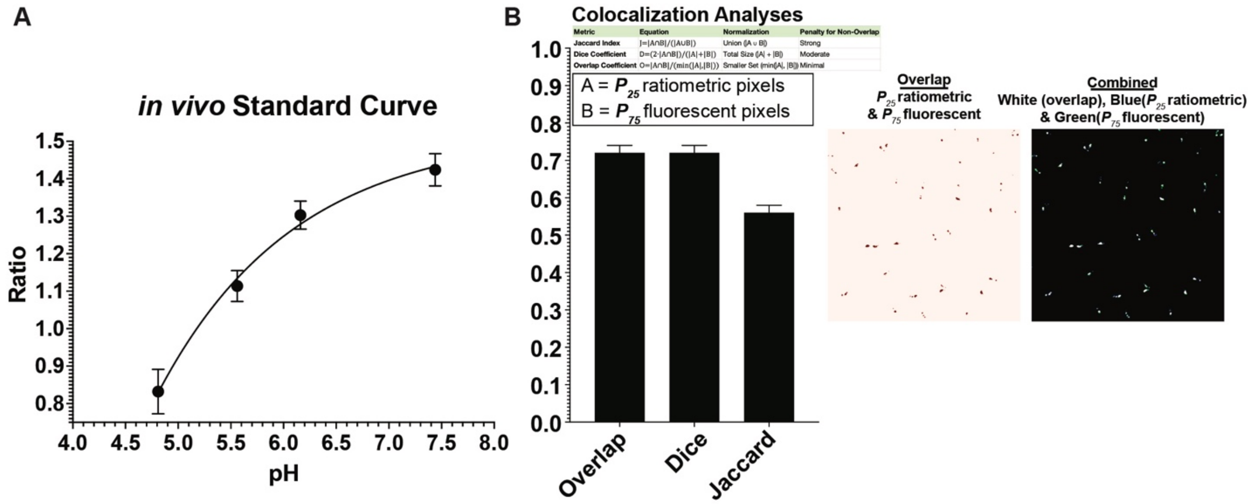


**Supplemental Figure 1.** (A) *In vivo* standard curve of RNase E-pHluorin2. (B) Colocalization analysis of *C. crescentus* expressing RNase E-pHluorin2, quantifying fractional overlap using the Jaccard index (0.72 ± 0.02), Dice coefficient (0.72 ± 0.2), and overlap coefficient (0.56 ± 0.02). Representative images highlight pixel overlap between the first quartile of ratiometric values (most acidic) and the fourth quartile of fluorescence intensity (most intense). Colocalization calculations were performed on composite images. Error bars represent standard deviations.

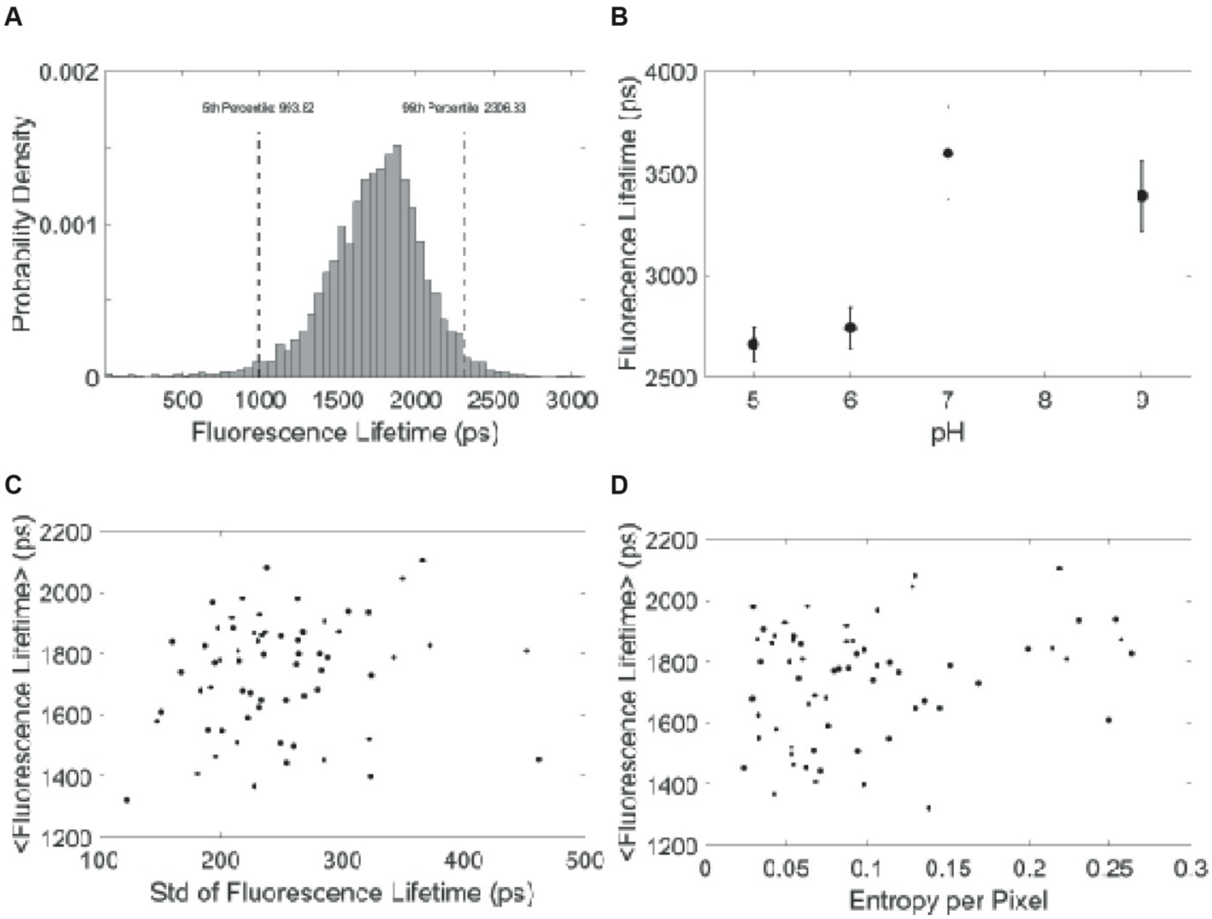


**Supplemental Figure 2.** (A) Probability distribution of fluorescence lifetimes acquired from pHluorin2-labeled RNase E in BR-body clusters of *Caulobacter crescentus* cells. (B) Average fluorescence lifetime of purified RNase E CTD-pHluorin2 in sodium acetate buffer (for pH 5.6) or Tris-Cl buffer (for pH 6.2, 7.5 and 9). Error bars represent standard deviations. (C) Scatter plot showing the average lifetime of each cluster and its standard deviation, indicating intra-cluster heterogeneity. (D) Scatter plot of the average lifetime of each cluster and corresponding entropy values calculated over 3-pixel grids, reflecting pixel-level heterogeneity within condensates.

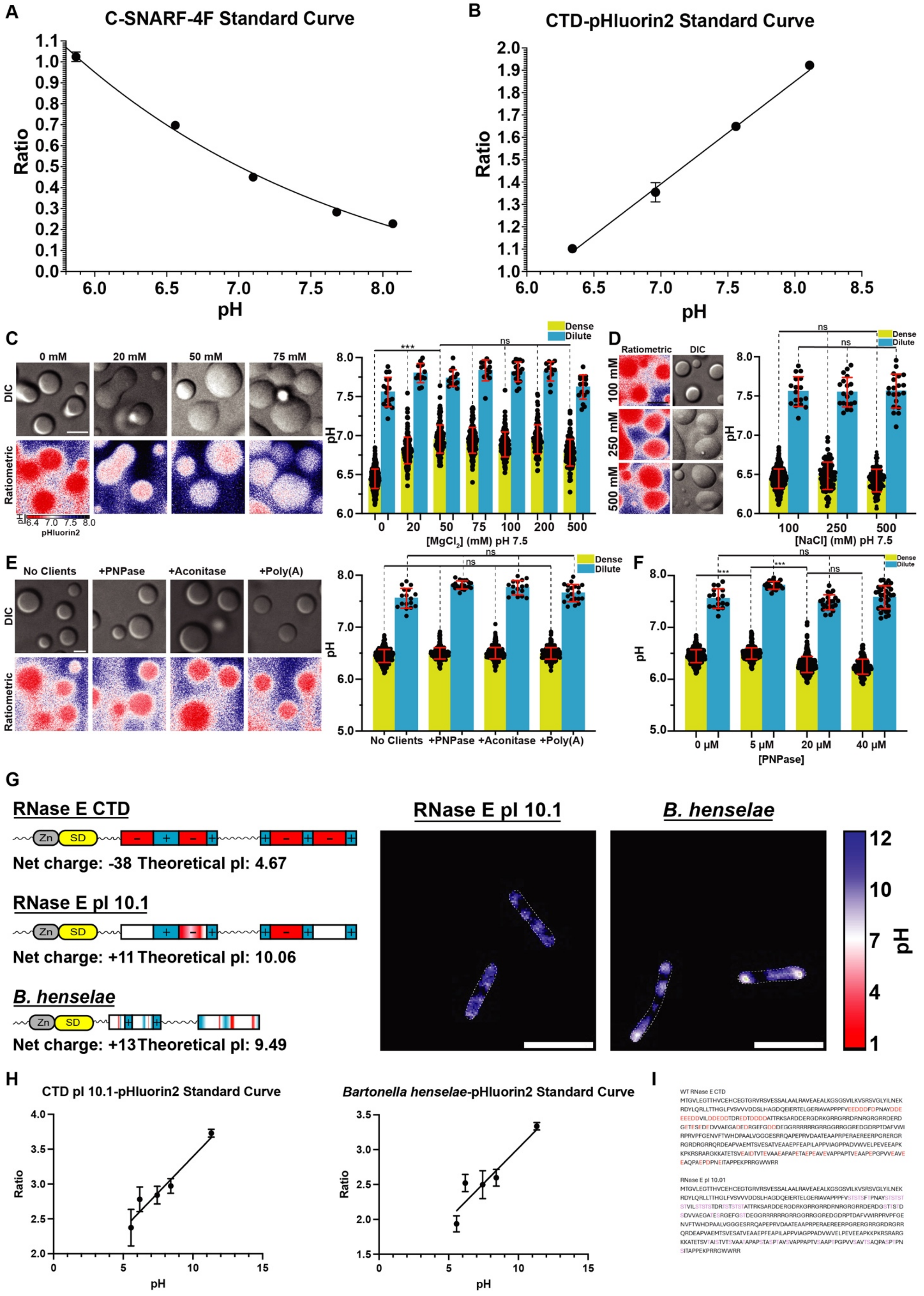
A
C-SNARF-4F Standard Curve
Ratio
pH
B
CTD-pHluorin2 Standard Curve
Ratio
pH
C
0 mM
20 mM
50 mM
75 mM
DIC
Ratiometric
pHluorin2
Dense
Dilute
[MgCl2] (mM) pH 7.5
D
Ratiometric
DIC
100 mM
250 mM
500 mM
[NaCl] (mM) pH 7.5
E
No Clients
+PNPase
+Aconitase
+Poly(A)
F
[PNPase]
G
RNase E CTD
Net charge: -38 Theoretical pI: 4.67
RNase E pI 10.1
Net charge: +11 Theoretical pI: 10.06
B. henselae
Net charge: +13 Theoretical pI: 9.49
RNase E pI 10.1
B. henselae
pH
H
CTD pI 10.1-pHluorin2 Standard Curve
Bartonella henselae-pHluorin2 Standard Curve
Ratio
pH
I
WT RNase E CTD
MTGVLEGTTHVCEHCEGTGRVRSVESSALAALRAVEAEALKGSGSVILKVSRSVGLYILNEK
RDYLQRLLTTHGLFVSVVVDDSLHAGDQEIERTELGERIAVAPPPFVEEDDDFDPNAYDDE
EEEDDVILDDEDDTDREDTDDDDATTRKSARDDERGDRKGRRGRRDRNRGRGRRDERD
GETESEDEDVVAEGADEDRGEFGDDDEGGRRRRRRGRRGGRRGGREDGDRPTDAFVWI
RPRVPFGENVFTWHDPAALVGGGESRRQAPEPRVDAATEAAPRPERAEREERPGRERGR
RGRDRGRRQRDEAPVAEMTSVESATVEAAEPFEAPILAPPVIAGPPADVWVELPEVEEAPK
KPKRSRARGKKATETSVEAIDTVTEVAAEAPAPETAEPEAVEVAPPAPTVEAAPEPGPVVEAVE
EAQPAEPDPNEITAPPEKPRRGWWRR
RNase E pI 10.01
MTGVLEGTTHVCEHCEGTGRVRSVESSALAALRAVEAEALKGSGSVILKVSRSVGLYILNEK
RDYLQRLLTTHGLFVSVVVDDSLHAGDQEIERTELGERIAVAPPPFVSTSTSFTPNAYSTSTST
STVILSTSTSTDRTSTSTSTATTRKSARDDERGDRKGRRGRRDRNRGRGRRDERDGSTTSTD
SDVVAEGATESRGEFGSTDEGGRRRRRRGRRGGRRGGREDGDRPTDAFVWIRPRVPFGE
NVFTWHDPAALVGGGESRRQAPEPRVDAATEAAPRPERAEREERPGRERGRRGRDRGRR
QRDEAPVAEMTSVESATVEAAEPFEAPILAPPVIAGPPADVWVELPEVEEAPKKPKRSRARG
KKATETSVTAISTVTSVAATAPAPSTASPTAVSVAPPAPTVSAAPTPGPVVSAVTSAQPASPTPN
SITAPPEKPRRGWWRR

**Supplemental Figure 3.** (A) *In vitro* calibration curve of C-SNARF-4F. (B) *In vitro* calibration curve of RNase E CTD-pHluorin2. (C) Representative images and bar graph illustrating the effect of titrating $MgCl_2$ into RNase E CTD-pHluorin2 condensates, showing a mild but consistent increase in dense-phase pH. (D) Representative images and bar graph depicting the effect of NaCl titration into RNase E CTD-pHluorin2 condensates (n=627), demonstrating negligible changes in dense-phase pH. (E) Ratiometric images of RNase E CTD-pHluorin2 following the addition of PNPase (n=253), aconitase (n=343), or poly(A) (n=283), each showing no significant impact on dense-phase pH. (F) Similarly, titrating PNPase up to 40 μM resulted in a minimally detectable pH change ($p<0.0001$). (G) Graphical representation of wild type RNase E CTD, a mutated version with pI = 10.1, and an RNase E CTD homolog from *Bartonella henselae* with pI = 9.5. Example ratiometric images of the RNase E CTD mutant and from *Bartonella henselae* are shown. (H) *In vivo* standard curves for the pHluorin2 chimeras of the RNase E CTD mutant and the *Bartonella henselae* homolog used for the analysis of ratiometric images are given. All error bars represent standard deviations. (I) WT RNase E CTD and the pI = 10.1 RNase E CTD mutant amino acid sequences with highlighted changes.

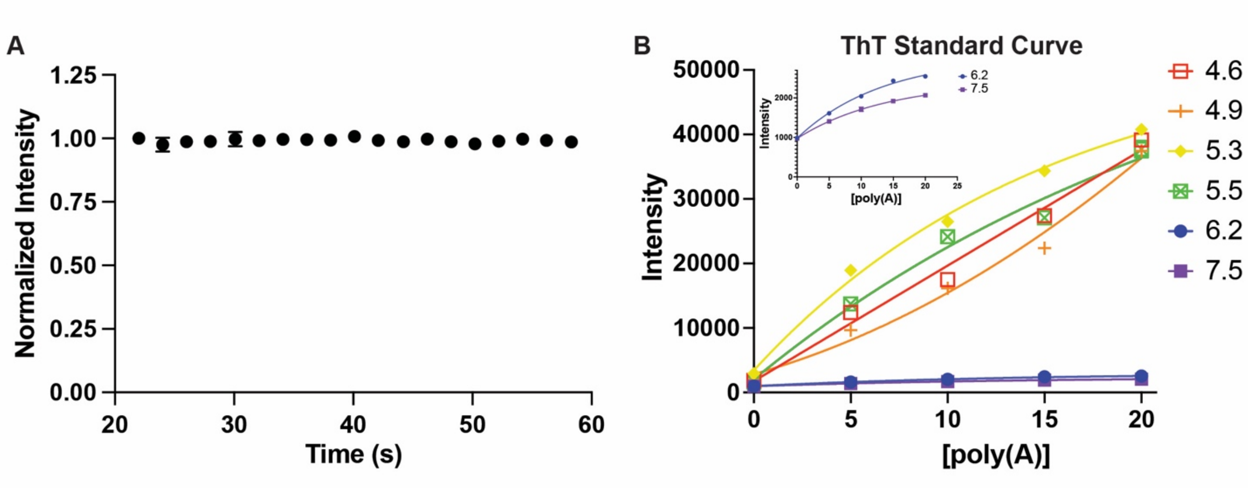


**Supplemental Figure 4.** (A) Normalized plot of Thioflavin T intensity when PNPase is inactive due to the lack of $Mg^{2+}$. (B) Thioflavin T intensity standard curves and fits as a function of poly(A) concentration. Each curve was generated based on the pH of the buffer used in each corresponding PNPase activity assay. The inset shows the dynamic response of Thioflavin T intensity at pH 6.2 and 7.5. All error bars represent standard deviations.

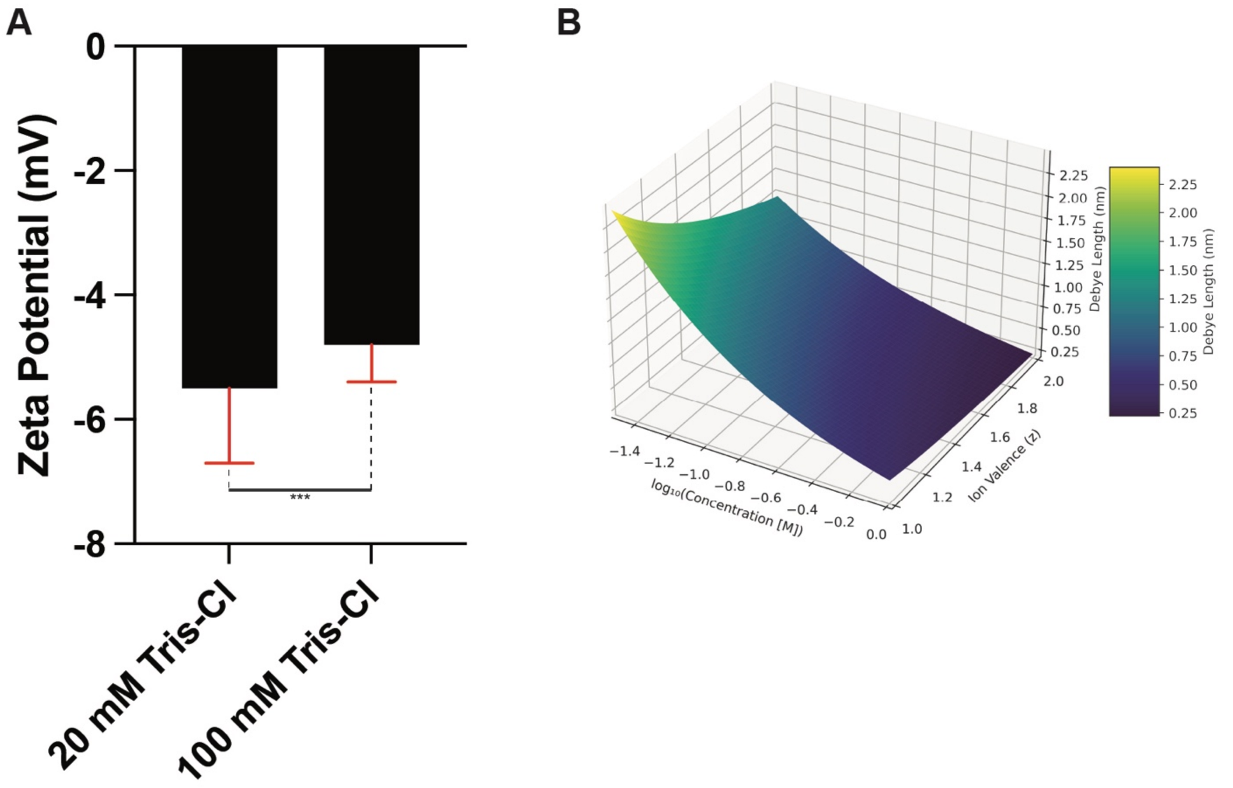


**Supplemental Figure 5.** (A) Zeta potential measurements under 5 mM Tris-Cl (pH gradient present) and 100 mM Tris-Cl (pH gradient not present) indicate that a non-zero zeta potential can be present without a pH gradient. (B) Plot of Debye length as a function of ion concentration and ion valence, illustrating its nonlinear dependence and sensitivity to charged species that may be present in either the dense or dilute phase of a two-phase system. All error bars represent standard deviations.
